# Methods in cancer research: Assessing therapy response of spheroid cultures by life cell imaging using a Cost-Effective Live-Dead Staining Protocol

**DOI:** 10.1101/2024.04.17.589882

**Authors:** Jaison Phour, Erik Vassella

## Abstract

Spheroid cultures of cancer cell lines or primary cells represent a more clinically relevant model for predicting therapy response compared to two-dimensional cell culture. However, current live-dead staining protocols used for treatment response in spheroid cultures are often expensive, toxic to the cells, or limited in their ability to monitor therapy response over an extended period due to reduced stability. In our study, we have developed a cost-effective method utilizing calcein-AM and Helix NP™ Blue for live-dead staining, enabling the monitoring of therapy response of spheroid cultures for up to 10 days. Using the example of glioblastoma cell lines and primary glioblastoma cells we show that spheroid cultures typically exhibit a green outer layer of viable cells, a turquoise mantle of hypoxic quiescent cells, and a blue core of necrotic cells when visualized using confocal microscopy. Upon treatment of spheroids with the alkylating agent temozolomide, we observed a reduction in the viability of glioblastoma cells after an incubation period of six to seven days. This method can also be adapted for monitoring therapy response in different cancer systems, offering a versatile and cost-effective approach for assessing therapy efficacy in three-dimensional culture models.

## Introduction

Precision oncology has become the standard of care of the treatment of different cancers harboring specific driver mutations. However, treatment response is often compromised due to intrinsic or acquired resistance mechanism in clinical practice. Genetic alterations, such as acquired mutations of drug targets, gain-of-function alterations of oncogenes in compensatory or bypass pathways, and epigenetic modifications play crucial roles in therapy resistance. Reduced uptake or increased efflux of the drug, enhanced DNA repair pathways, activation of survival pathways, tumor cell plasticity, and modulation of the tumor microenvironment are additional mechanisms contributing to treatment response [1,2].

Indeed, the ineffectiveness of drug treatment is responsible for up to 90% of all cancer-related deaths [3,4].

Three-dimensional (3D) spheroid cultures of cancer cell lines or primary cells represent a more clinically relevant model for predicting therapy response compared to two-dimensional cell culture as they share some features of tumor tissues (Fig. 1). Notably, factors such as hypoxia, acidosis, and cell-to-cell interactions, which contribute to treatment response, are better reflected in 3D cultures. Consequently, the sensitivity to drugs is often similar in spheroids and tumor tissues, whereas 2D models are a less reliable predictor of therapy response [5,6].

**Figure 1:**
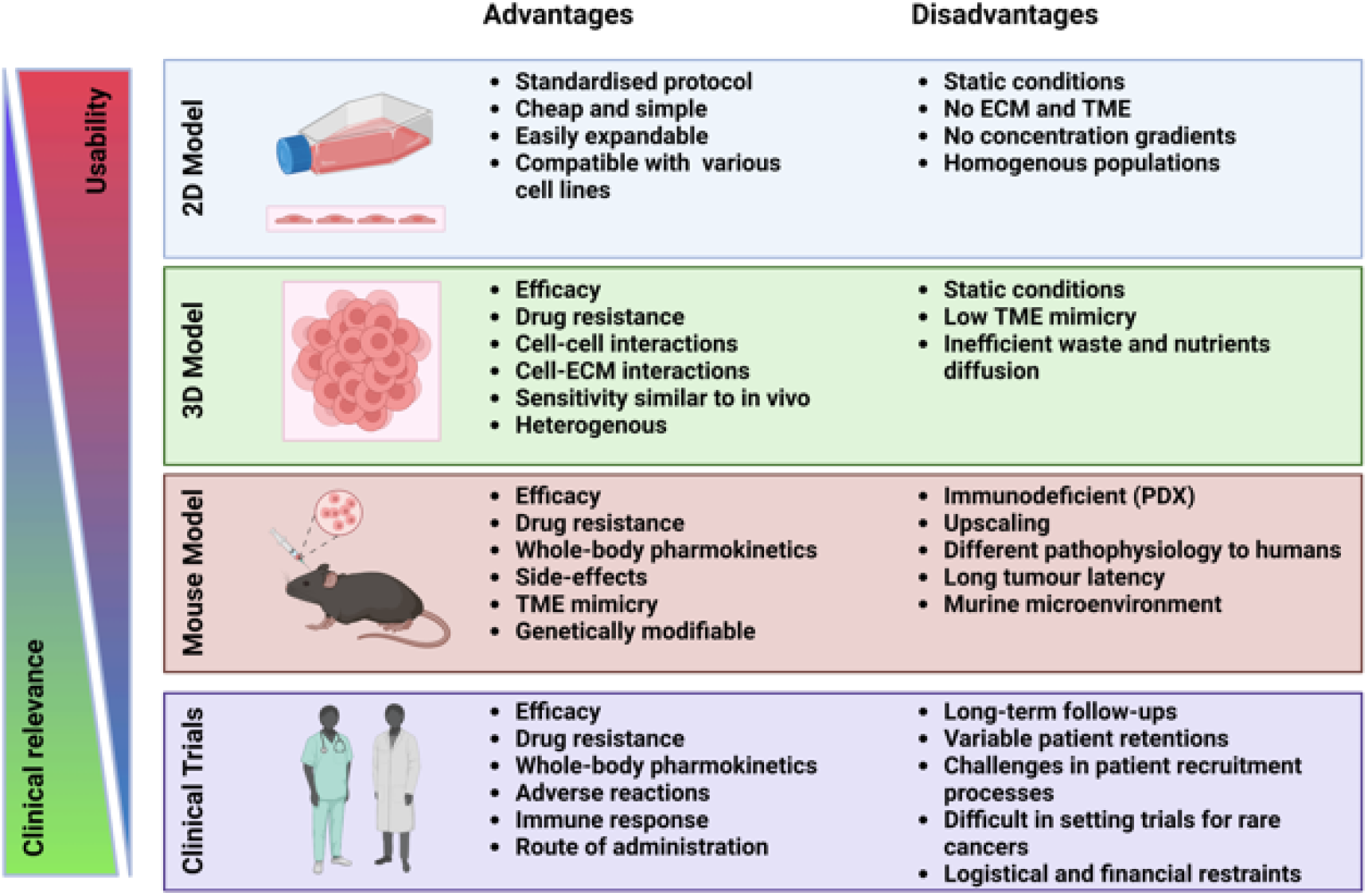
Pre-clinical models. Advantages and disadvantages of pre-clinical models for predicting drug response in patients. Adapted from Law et al., 2021. Created with BioRender.com.

Current live-dead staining protocols used for assessing treatment response are often expensive, toxic to the cells, or limited in their ability to monitor therapy response over an extended period due to reduced stability. To address this issue, we present a cost-effective method utilizing calcein-AM and Helix NP™ Blue for live-dead staining and a protocol for live imaging using the mica microhub imaging system (Leica, v.1.0.1) and Leica application suit X (LAS X) v.6.2.2.28360, enabling the monitoring of therapy response of spheroid cultures for up to 10 days.

## Materials and Methods

### Cell lines and culture conditions

The glioblastoma cancer cell lines LN-18 (ATCC CRL-2610™) and LN-229 (ATCC CRL-2611™) and primary glioblastoma stem cells intraoperatively collected from a patient with glioblastoma [7] were used in this study. Cell lines were genotyped and authenticated by Microsynth (Switzerland) and tested negative for mycoplasma infections. They were cultivated in DMEM, low glucose (Sigma-Aldrich) supplemented with 5% FBS (Sigma-Aldrich) and 2mM L-Glutamine (Sigma-Aldrich). Primary glioma stem cells (GSCs) were cultured in DMEM/F12 (ThermoFisher) supplemented with 1× B27, 20 ng/ml epidermal growth factor (EGF), and 20 ng/ml basic fibroblast growth factor (bFGF). If not otherwise stated cells were treated with 100_μ_M temozolomide or vehicle. LN-18 and GSC4 were also treated with 10_μ_M O6-Benzylguanine (O6BG) to enhance temozolomide response.

### Spheroid assembly

Monolayer cultures of cell lines were allowed to reach maximal confluency of 70% before initiating spheroid culture. The viability and cell number of the cell suspension used for spheroid culture were verified by trypan blue staining. Only cell cultures showing a cell viability of at least 90% were used for spheroid culture. For spheroid formation, 2x 104 vital cells per well were seeded in 100 μl of medium in a 96-well BIOFLOAT™ round bottom plate (Sarstedt). Spheroids were formed by centrifugation of the 96-well plate for 10 min at 500 g. Subsequently, spheroids were allowed to form by incubation at 37°C, 5% CO2 in a humid chamber for 4 days. Temozolomide was then added at final concentration of 100μM, and incubation was continued for up to seven days.

### Live and dead staining

At 96 hours post-seeding, the culture medium was supplemented with 0.1 mM Copper(II) sulfate (CuSO4) (Sigma-Aldrich), 2 μM calcein-acetoxymethyl (calcein-AM) (BioLegend), and 2.5 μM Helix NP™ Blue (BioLegend), along with 100 μM temozolomide or vehicle. Every 3 to 4 days, 50 μl of culture supernatant was replenished with fresh reagents.

### Determination of cell viability following treatment

The viability of treated tumor spheroids was assessed using the mica microhub imaging system (Leica) through live-cell imaging with confocal microscopy on day 0 and day 7. Imaging was performed within the first 30min after staining. ImageJ v.1.53t (NIH) software and Fiji (v. 1.54i) was utilized to generate Z-stack images using the Z-Projection function with average intensity, and the ICY BioImage Analysis Tool v.2.5.2.0 (Institute Pasteur & France-BioImaging) was employed for active contour detection (https://gitlab.pasteur.fr/bia/active-contour) and spot detection (https://gitlab.pasteur.fr/bia/spot-detector) of fluorescent signals. The UnDecimated Wavelet Transform detector (UDWTWaveletDetector) algorithm, which detects bright spots over dark background, along with additional SizeFiltering, was used in ICY to detect spots corresponding to cells. Survival rates were further calculated as followed.

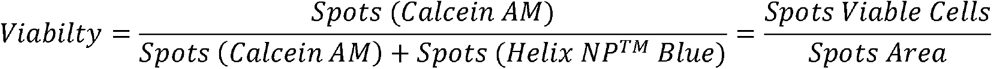

A detailed description of the protocol is attached as a supplementary file.

### Statistical analysis

**GraphPad Prism software (version 9.4.1) was employed for statistical analysis. An ordinary one-way ANOVA with Šídák’s multiple comparisons test was conducted to assess differences between the control group and the treated groups. The data are presented as the mean ± standard deviation (SD).Results**

To evaluate therapy response, we developed a protocol for spheroid culture formation and conducted life-dead imaging using the mica microhub imaging system (Leica) and ICY BioImage Analysis Tool using spot detector plugin. Our approach involves seeding cells in BIOFLOAT™ plates and collecting them by centrifugation, which promotes the formation of spheroids due to the poor attachment of cells to the plate surface. The use of U-profile round bottom plates ensures that typically only one spheroid per well is formed, although the ability to generate perfectly shaped spheroids may vary depending on the cell type and culture conditions. For example, we observed that LN-229 cells form perfectly shaped spherical structures, while LN-18 cells form non-uniform aggregates. Even primary GSC4 cells, which grow in suspension, can be used to generate spheroids. These spheroids exhibit characteristics resembling glioblastoma tumors, including an outer layer of proliferating cells, a mantle of hypoxic-quiescent cells, and a necrotic core. (Fig. 2A). Although modifications to the cell culture medium, such as adjusting the concentration of bovine serum or supplementing with conditioned medium, may improve spheroid formation, our experiments showed that using ultra-low attachment plates did not yield significant improvements.

**Figure 2:**
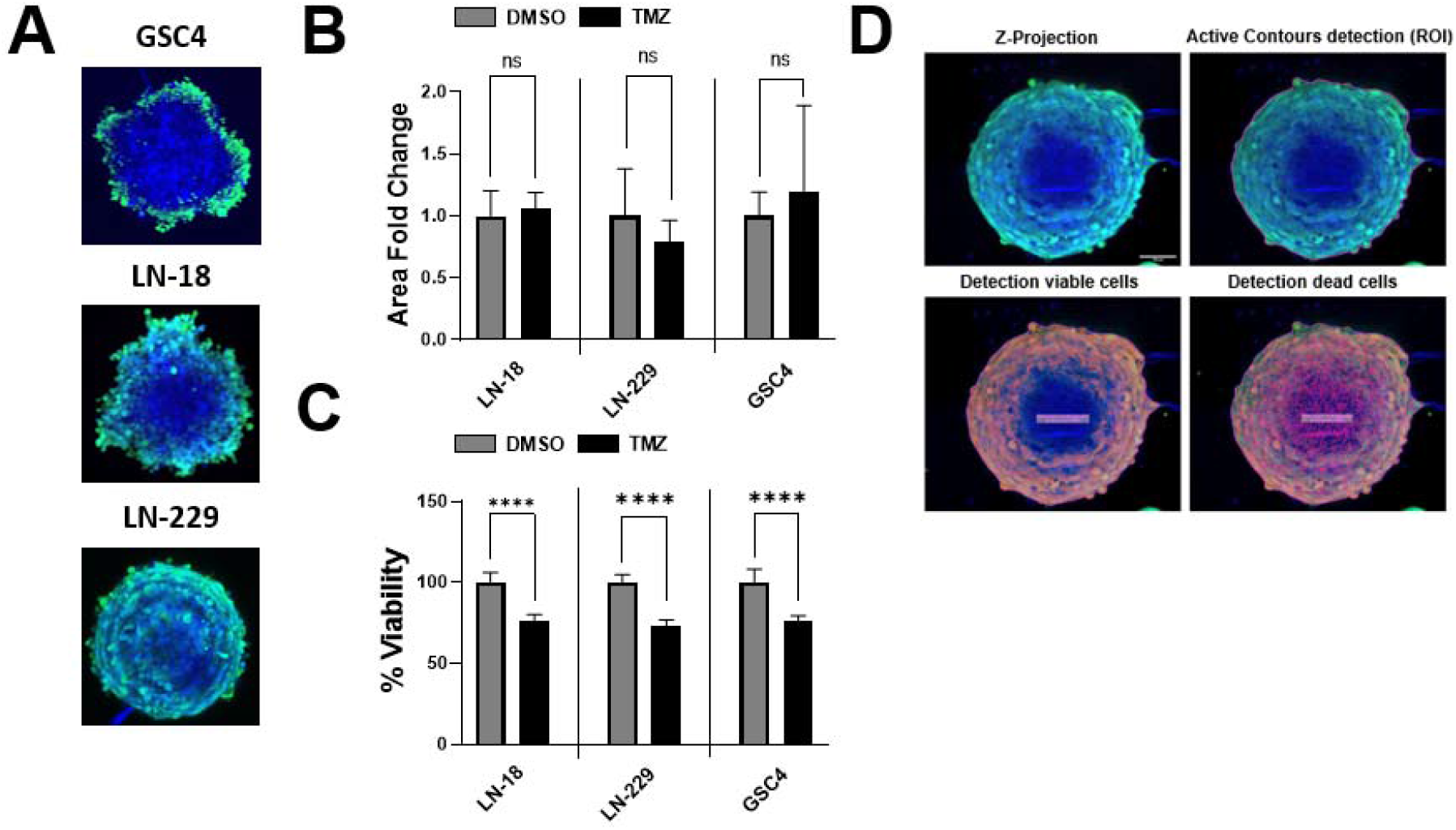
Spheroid model. (A) Representative images of GSC4, LN-18 and LN-229 spheroids cultured in the absence of a drug for 7 days. Viable proliferative cells of the outer layer are shown in green, hypoxic-quiescence cells from the mantle are shown in turquoise and necrotic cells from the core are shown in blue. (B) Spheroid size and (C) percent viable cells relative to total cell counts normalized to day 0 in the presence of temozolomide (TMZ) or DMSO control. (D) ICY BioImage Analysis Tool utilizing spot detection plugin. This detection method of Z-Protection (upper left) involves determining regions of interest (upper right), detection of viable cells (lower left) and detection of dead cells (lower right).

For evaluating therapy response based on spheroid size, the formation of perfectly shaped spheroids is crucial. The criteria used in the literature, which typically involves visual inspection of the spheroid size, are often arbitrary [8–10]. Spheroid size has a significant impact on experimental outcomes, with larger spheroids potentially showing no difference in size following treatment with drugs like temozolomide (Fig. 2B) [11]. While previous studies have addressed the issue of mass transport limitation in larger spheroids hindering drug responsiveness, our observations suggest that relying solely on size or area measurements may not adequately detect drug response in tumor spheroids. Instead, we propose that in larger spheroids, dead cells may be less decomposed and remain as part of the spheroid, especially in experiments lasting more than 7 days (Fig. 2C). For the viability assessment, we initially generate a Z-Projection with average intensity using ImageJ/Fiji. Subsequently, we mark our region of interest (ROI) and select viable and dead cells through spot detection using ICY BioImage Analysis Tool (Fig. 2D).

## Discussion

Our results demonstrate that in cases where size or area measurements fail to detect drug response, live-dead cell viability staining can still accurately assess the drug response. Here, we describe a staining protocol using calcein-AM and Helix NP™ Blue (also known as Sytox™ Blue) for live/dead imaging. Calcein-AM is membrane-permeable and is cleaved by intracellular esterases, producing calcein, a green-fluorescent product, which accumulates in the cytoplasm of living cells [12]. The addition of CuSO4 to the culture medium is crucial as it effectively reduces background staining derived from lysed calcein-positive cells [13]. Helix NP™ Blue is impermeant to live cells and binds to nucleic acids of dead cells emitting a blue fluorescent light. Thus, this protocol is effective for live-dead cell discrimination.

This protocol is cost-effective and does not cause any cytotoxic effects. The fluorescent signal remains stable for up to 4 days. For longer incubation periods, we recommend replenishing the culture medium with fresh reagents before analysis. Fluorescence intensity may fluctuate due to fading of the fluorescent signal over time and may increase again after replenishing with fresh reagents. However, based on our experience, fluorescence intensity typically remains above the thresholds that we set for live and dead staining. Consequently, fluctuations in fluorescence intensity do not influence the proportion of live and dead cells. To correct for the proportion of live and dead cells resulting from drug treatment, we recommend normalizing the proportion of live and dead cells after drug exposure to the proportion observed at the beginning of the experiment, prior to drug administration (day 0). This approach enables us to evaluate acurate therapy responses over time and facilitates comparisons between spheroids generated from cells with different genetic backgrounds. Our method utilizes spot detection plugin in ICY BioImage Analysis Tool for determining the number of live and dead cells, enabling accurate measurements of viability of the individual tumor spheroid during time-course experiments. In contrast, Bulin et al., utilizes fluorescent intensity of calcein-AM and propidium iodide (PI) stain to calculate viability [14]. One drawback of using propidium iodide (PI) for staining is its cytotoxic effect, limiting its application to endpoint measurements. In summary, calcein-AM staining in combination with Helix NP™ Blue staining offers the advantage of evaluating both parameters during time-course experiments. To our knowledge, this combination of live and dead cell staining has not yet been utilized.

The proposed method also allows for the comparison of therapy responses between two populations within the same spheroid. To achieve this, both populations can be transduced with different fluorescent proteins or pre-labelled with different dyes. However, it is important to choose additional fluorescent proteins or dyes that do not overlap in the emission spectrum of live and dead stains.

A more accurate method for detecting viability would involve identifying specific cells in a 3D-rendered format rather than detecting spots in Z-projections. This approach would yield a more precise percentage of viable cells. Achieving this requires an increased number of layers in the Z-stack to optimize 3D deconvolution and rendering for cell detection. However, this alternative protocol is highly time-consuming and is only recommended for small series of experiments. To facilitate large-scale screenings, generating widefield images using the mica microhub imaging system (Leica) may be an alternative to confocal images, as it allows for faster processing of data. However, in our experience, widefield images have lower quality and do not produce reliable results for longer incubation periods.

## Conclusions

Assessing therapy responses using spheroid cultures represents a more clinically relevant model compared to two-dimensional cell culture. However, live-dead staining provides a more accurate means of assessing therapy response compared to measuring spheroid size. We have developed a robust and cost-effective protocol for live-dead staining and analysis using confocal microscopy, which is suitable for time-course experiments over an extended period. This method can also be adapted for use with more complex systems such as tumoroids or organotypic cultures of patient-derived tissue explants. Furthermore, derivates of calcein-AM and Helix NP™ Blue with different fluorescent spectra exist, enabling a broader range of applications and making them adaptable for multicolor staining of spatial analysis. Live cell imaging, as conducted by the mica microhub imaging system (Leica), can also be applied to other commercially available widefield or confocal fluorescence microscopy systems used in other laboratories.

## Disclosure

### Author Contributions

Design and direction of experiments, conceptualization, methodology, analysis of data J.P.; writing manuscript – original draft preparation J.P.; review and editing manuscript E.V.; supervision E.V.

