## Supplementary file for "Methods in cancer research: Assessing therapy response of spheroid cultures by life cell imaging using a Cost-Effective Live-Dead Staining Protocol"

### Detailed description of the used analysis softwares

**LAS X** (v.6.2.2.28360)

Excitation and emission spectra for Helix NP™ Blue and calcein AM were downloaded from fpbase.org/spectra and installed on the Leica application suit X (LAS X) software to image the spheroids on mica microhub imaging system v.1.0.1 (Leica). We utilized 20x magnification images for our analysis. The software automatically set the illumination intensity, which was configured to confocal grade and high quality on day 0. The same illumination settings were consistently applied to all other time points. For each imaging step, a focus map was generated with a focus point for each spheroid, and a z-stack with a Z-Step size of 30 µm was captured. Raw images were subsequently exported as TIFF files, preserving all raw information and channels.

**ImageJ** (v.1.53t) / **Fiji** (v. 1.54i)

The raw images were imported to ImageJ, and the channels were merged by color. Z-projection of the images were performed using average intensity, and the results were saved as TIFF-files.

**ICY BioImage Analysis Tool** (v.2.5.2.0)

The Z-stack images were imported, and the Active Contour plugin (https://gitlab.pasteur.fr/bia/active-contour) was utilized for segmentation of the entire spheroid. Briefly, a 2D ROI was drawn around the object of interest, and parameters were adjusted accordingly. Since we aimed to detect the whole spheroid as the ROI, we set the edge weight to 1 and region weight to 0 on the green channel, expecting a viable outer layer. Additionally, the evolution time-step was adjusted based on the background to expedite the process, while default settings were maintained for the other parameters. A detailed user documentation for this process is available on the official Active Contour documentation page on the ICY website (http://icy.bioimageanalysis.org/plugin/active-contours/).

To segment the cells within the spheroids, we utilized the spot detector plugin (https://gitlab.pasteur.fr/bia/spot-detector). This plugin detects spots, which can represent nuclei or cells depending on the objective. It employs the UnDecimated Wavelet Transform detector (UDWTWaveletDetector) algorithm, designed to detect spots, even in the presence of high image noise. We configured the plugin to detect brighter spots on dark backgrounds and adjusted the spot size based on the cell line. The pixel size was manually measured for a single cell to select the appropriate scale, which is cell line dependent. Size criteria were also included in the filtering box to further reduce background noise. Additional details about this plugin can be found on the ICY websity (http://icy.bioimageanalysis.org/plugin/spot-detector/).

Once all parameters were set, spots were detected for each channel, and viability was computed as followed.

$$Viabilty = \frac{Spots\ (Calcein\ AM)}{Spots\ (Calcein\ AM) + Spots\ (Helix\ NP\ ^{TM} Blue)} = \frac{Spots\ Viable\ Cells}{Spots\ Area}$$

To correct for the proportion of live and dead cells resulting from drug treatment, we normalized the proportion of live and dead cells after drug exposure (day 7) to the proportion observed at the beginning of the experiment, prior to drug administration (day 0).

$$Viabilty_{Normalized} = \frac{Viabilty\ _{Day7}}{Viabilty\ _{Day0}}$$
